# Sensitivity analysis-guided Bayesian Parameter Estimation for Neural Mass Models: Applications in Epilepsy

**DOI:** 10.1101/2023.04.03.535345

**Authors:** Narayan Puthanmadam Subramaniyam, Jari Hyttinen

**Affiliations:** Faculty of Medicine and Health Technology, Tampere University, Tampere 33520, Finland

## Abstract

It is well established that neural mass models (NMMs) can effectively simulate the mesoscopic and macroscopic dynamics of electroencephalography (EEG), including epileptic EEG. However NMMs are characterized by a high-dimensional parameter space and a lack of knowledge on what NMM parameters can be reliably estimated, thus limiting their application to clinical EEG data. In this article, we analyze the parameter sensitivity of Jansen and Rit NMM (JR NMM) in order to identify the most sensitive NMM parameters for reliable parameter estimation from EEG data. We also propose a joint estimation method for NMM states and parameters based on expectation–maximization combined with unscented Kalman smoother (UKS-EM). Global sensitivity analysis methods including Morris method and Sobol method are used to perform sensitivity analysis. Results from both the Morris and Sobol method show that the average inhibitory synaptic gain, *B* and the time constant of the average inhibitory post-synaptic potentials (PSPs), *b*^−1^ have significant impact on the JR NMM output along with having the least interaction with other model parameters. The UKS-EM method for estimating the parameters *B* and *b* is validated using simulations under varying levels of measurement noise. Finally we apply the UKS-EM algorithm to intracranial EEG data from 16 epileptic patients. Our results, both at individual and group-level show that the parameters *B* and *b* change significantly between the pre-seizure and seizure period, and between the seizure and post-seizure period, with transition to seizure characterized by decrease in average *B* and high frequency activity in seizure characterized by an increase in *b*. These results establish sensitivity analysis guided Bayesian parameter estimation as a powerful tool for reducing the parameter space of high dimensional NMMs enabling reliable and efficient estimation of the most sensitive NMM parameters, with the potential for online and fast tracking of NMM parameters in applications such as seizure tracking and control.

## 1 Introduction

Epilepsy is characterized by epileptic seizures, which are recurrent and unpredictable interruptions of the normal brain function [1]. Despite the development of more than 20 anti-epileptic drugs, around 30% of the ≈65 million epileptic patients remain pharmacoresistant [2]. Interictal spikes, which occur between the seizures, are synchronous neuronal discharges and have shown to last tens to hundreds of milliseconds as revealed by electroencephalographic (EEG) recordings from hippocampal structures [3]. Ictal events, or seizures, last for several tens of seconds and are characterized by rhythmic discharges [4]. The process leading to the seizure onset is defined as ictogenesis and the biological mechanisms underlying ictogenesis remain poorly understood [4].

Computational models in neuroscience are useful to explain experimental findings and generate new testable hypotheses [5]. It is well established that mean-field approximation models such as neural mass models (NMMs) are suited to study the mesoscopic and macroscopic dynamics of EEG [6, 7]. NMMs describe the interaction between the pyramidal neurons, excitatory interneurons and inhibitory interneurons using a system of nonlinear ordinary differential equations (ODEs). Although NMMs represent the aggregated activity of neuronal populations rather than a single neuron dynamics, the aggregated parameters of NMMs have biological significance. Several studies have shown that various kinds of EEG activity observed in epileptic patients can be simulated by tuning the parameters of NMMs related to the average excitatory and inhibitory synaptic gains [8, 9]. Although there are several NMMs proposed in the literature, the Jansen and Rit (JR) model [8] is the most commonly used NMM describing the interactions between neuronal populations using a set of six nonlinear ODEs and can produce a variety of EEG dynamics including normal EEG, EEG with sporadic spikes, sustained rhythmic discharges and quasi-sinusoidal activity [9].

Knowledge on how various parameters of NMMs contribute in producing different types of epileptic EEG activity can provide new insights on the mechanism underlying epilepsy. Furthermore, identifying such parameter changes from EEG recordings can aid in the treatment of epilepsy from the perspective of designing devices that are, for example, able to deliver electrical stimulation on demand to reduce or eventually eliminate seizure activity [10]. Although various approaches for estimating parameters of NMMs have been proposed [9, 11] approaches based on sigma-point Kalman filters [12, 13, 14, 15] have been widely used [10, 16, 17, 18] as they are considered as better alternatives to extended Kalman filters when dealing with highly nonlinear systems [13, 12] such as NMMs.

Since the effect of the excitatory and inhibitory gain parameters on the NMM model output has been extensively studied [9], previous studies on parameter estimation of NMMs based on sigma-point Kalman filters have mostly focused on estimating changes related to these parameters during transition to seizure, assuming all the other NMM parameters to be constant [10, 16, 19]. However, NMMs typically contain several parameters, with certain parameters influencing the model output more than the others. Also, certain parameters might have large variations between different conditions (for example ictal and interictal conditions) while other parameters might have little to no effect. Furthermore, some of these parameters may be highly correlated (nonlinearly) and thus change in one parameter can be compensated by a change in another parameter. As seen in Table 1, even a simple JR NMM has 13 parameters. How the variations in these parameters affect the output of the JR NMM and what is the degree of interactions between these parameters and which of these parameters can be reliably estimated using sigma-point Kalman filters from EEG data has not been investigated so far.

**Table 1:** The Jansen-Rit NMM parameters and their interpretation. The NMM produces normal EEG as its output when the parameters are fixed to these values [5].

| Parameter | Interpretation | Value |
| --- | --- | --- |
| $A$ | Average excitatory synaptic gain | 6 mV |
| $B$ | Average inhibitory synaptic gain | 70 mV |
| $1/a$ | Time constant of average excitatory PSPs | $a = 100 \text{ s}^{-1}$ |
| $1/b$ | Time constant of average inhibitory PSPs | $b = 50 \text{ s}^{-1}$ |
| $C_1, C_2$ | Average number of synaptic contacts in the feedback excitatory loop | $C_1 = 135, C_2 = 108$ |
| $C_3, C_4$ | Average number of synaptic contacts in the slow feedback inhibitory loop | $C_3 = 33.75, C_4 = 33.75$ |
| $v_0, e_0, r$ | Parameters describing the Sigmoid function $\mathcal{S}$ | $v_0 = 6 \text{ mV}, e_0 = 2.5 \text{ s}^{-1},$<br>$r = 0.56 \text{ mV}^{-1}$ |
| $p(t) \sim \mathcal{N}(\mu, \sigma)$ | Input noise | $\mu = 90, \sigma = 30$ |

Sensitivity analysis (SA) is a widely used technique to understand how changes in model parameters affect the model output [20]. SA can provide useful insight on how sensitive the model output is to its parameters by identifying which of the model parameters have the most influence on the model output and to which parameters the model output is insensitive. Thus, SA can also be used as a tool for model simplification as values of the insensitive parameters can be assumed to be fixed for various conditions. Furthermore, since SA can identify which model parameters contribute the most to the changes in model output, these sensitive model parameters can be considered as the targets for parameter estimation from experimental data. SA has been applied in various fields including environmental modeling [21], chemical kinetics [22, 23], biomedical sciences [24] and economic modeling [25] to mention a few.

SA can be broadly categorized into local and global SA. In local SA, the impact of small changes to the input parameter on the model output is studied [26] and only one parameter at a time is varied while all other parameters fixed at their reference value. Partial derivatives (analytical or numerical) of the model output with respect to the individual parameters are obtained analytically or approximated numerically, to determine how changes in one parameter impacts the output. However local SA methods are limited in their usefulness as the results of this method are only valid near a local point [27, 26].

In contrast, global SA can be used to investigate the sensitivities of model outputs when there can be large variations in the parameter values [28]. Global SA method allows to explore range of parameter values using sampling strategies. Several studies have shown that simple random sampling strategies, although an easy method to generate parameter samples require a large number of samples to cover the entire range of the model input parameters and thus are computationally demanding [24]. In contrast, Latin hypercube sampling (LHS) strategy proposed by [29] and later extended by [30] is a more efficient method to generate random samples that sufficiently cover the parameter space using fewer sampling iterations compared to random sampling [31, 24]. LHS is thus more commonly used when applying global SA methods.

In this work we apply two global SA techniques – the Morris method [32, 33] and variance-based Sobol method [20] to identify the model parameters that most influence the JR NMM output, i.e. the EEG data. Then we propose a parameter estimation technique using an expectation–maximization (EM) algorithm with unscented Kalman filter (UKF) and Unscented Rauch-Tung-Striebel smoother(also known as unscented Kalman smoother (UKS)) [34], hereafter referred to as UKS-EM algorithm, to identify the most influential parameters of the JR NMM. The UKF belongs to the family of sigma-point Kalman filters [13] and is based on unscented transformation for calculating the statistics of a random variable undergoing a nonlinear transformation [13, 12]. We resort to UKS-EM algorithm rather than using just UKF is because, in the UKF approach, the noise covariance matrices have to be tuned to improve the state and parameter estimation and this requires prior knowledge. Within the UKS-EM framework, we are able to derive the maximum likelihood estimates for the noise covariance matrices, avoiding the need to have prior knowledge on statistical characteristics of system noises to tune the covariance matrices optimally [35]. We validate the performance of our parameter estimation method on simulated EEG data generated by varying the most influential parameters of JR NMM, based on the results from SA under varying levels of measurement noise. Finally, to demonstrate the validity of our approach, we used intracranial EEG (iEEG) data from 16 epileptic patients obtained from an open database and tracked the changes in the influential JR NMM parameters with UKS-EM algorithm during the pre-seizure, seizure and post-seizure period.

## 2 Neural mass model

The post-synaptic potential of a population of neurons *n,* can be given as,

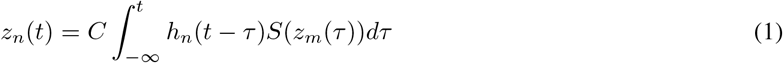

where *C* is the average number of synaptic contacts between populations *n* and *m* (for example between pyramidal neurons and inhibitory interneurons). Here, *h_n_*(·) is the synaptic impulse response function (Green’s function) and is given as,

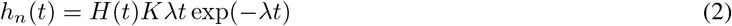

where *H*(·) is a Heaviside step function, *λ* is the response time constant and *K* represents the average synaptic gain for the neuronal population *n*. In equation 1, 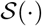 is often modeled as a logistic function and is given as,

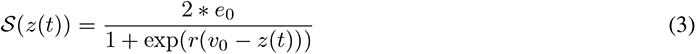

where 2 * *e*_0_ is the maximum value of the function, *r* represents the steepness of the curve and *v*_0_ represents the excitability threshold.

The convolution given in equation 1 can be considered as a solution to the initial value problem 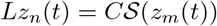, where the linear differential operator *L* is given as,

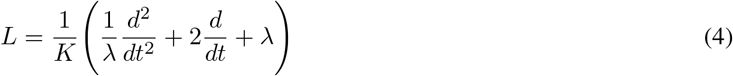

Thus, we get

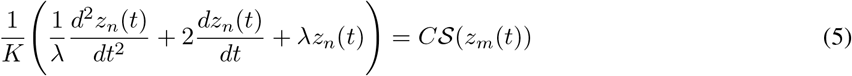

Re-arranging, we have

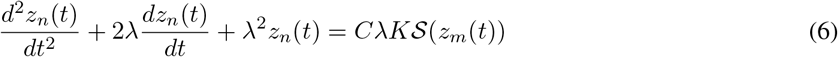

Switching to Newton’s notation for differentiation and re-writing the second-order ODE as two coupled first-order ODEs we have,

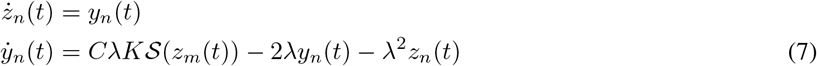

Equation 7 now represents an interaction between two neuronal population *n* and *m*.

### 2.1 Jansen-Rit NMM

The Jansen-Rit NMM models the interaction between population of pyramidal neurons and inhibitory interneurons. Two subsets of pyramidal neurons are considered: primary and secondary pyramidal neurons. The secondary population of pyramidal neurons represent distant neurons that have synaptic connections with the primary pyramidal population. The primary pyramidal neurons receives excitatory inputs from secondary pyramidal neurons (collateral excitation) and inhibitory input from inhibitory interneurons. Both inhibitory interneurons and secondary pyramidal neurons receive excitatory input from pyramidal interneurons.

These excitatory and inhibitory interactions are modeled via the following set of nonlinear ODEs which take the general form shown in Equation 7,

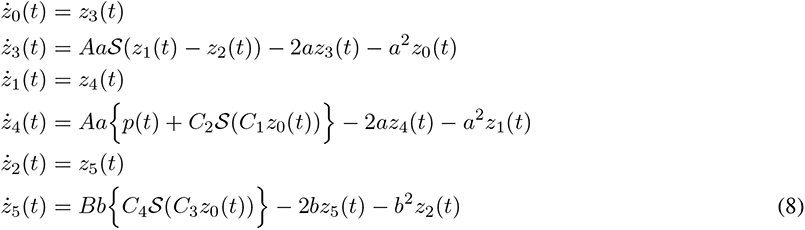

The output modelling the EEG activity is given as,

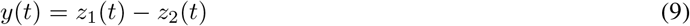

where *z*_1_(*t*) and *z*_2_(*t*) are the excitatory and inhibitory post-synaptic potentials.

### 2.2 Discrete-time state space representation

We can also describe the continuous-time, nonlinear dynamical system shown in equation 8 and 9 as a discrete-time nonlinear state-space model,

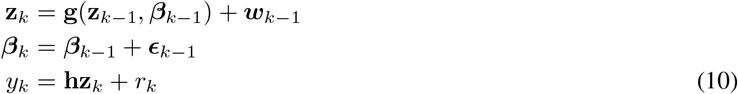

where 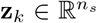 represents the unobserved state, 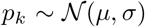 represents the Gaussian noise input and 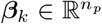 represents the unknown parameters of NMM at time step *k* and **h** is the observation vector that models equation 9. The observed (single channel) EEG measurement signal is represented as *y_k_*. The function **g** describes the discrete-time version of the nonlinear function described by the ODEs shown in equation 8.

The terms *w_k_, ϵ_k_* and *r_k_* represent model errors and are assumed to be Gaussian, i.e. 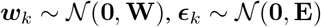 and 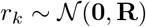 respectively. Furthermore, the state **z***_k_*, parameter ***β**_k_* and observation vector **h** is given as,

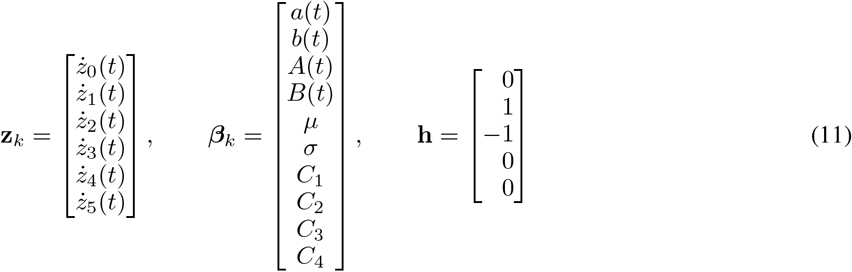

## 3 Sensitivity analysis of the Jansen-Rit NMM

We use two global SA methods, – the Morris method and Sobol’s indices to investigate the impact of input parameters on the model output. LHS strategy is used in both these methods to sample a range of parameter values.

Let *f*(·) represent some mathematical model and ***β*** = (*β*_1_, *β*_2_,…, *β_n_p__*) represent vector of *n_p_* input parameters. The model output y can be represented as,

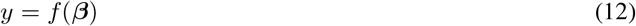

In case of the Jansen-Rit NMM, the model input parameter vector ***β*** is given by equation 11 and the (scalar) model output *y* represents the EEG defined in equation 9.

### 3.1 Morris method

One of the commonly used global SA methods is the Morris method [32] and it can be used to rank the importance of model parameters qualitatively. The Morris method is particularly useful when there are many input variables describing a system or a model and it is not practical to analyze all of them at once.

The Morris method is based on randomized one-at-a-time (OAT) design, which is based on sampling the partial derivatives of the output with respect to a one input parameter when all the other parameters are fixed. For the Jansen-Rit NMM described in Section 2.2, the elementary effect (EE) of an *i*–th parameter in the parameter vector ***β*** on the model output is given as,

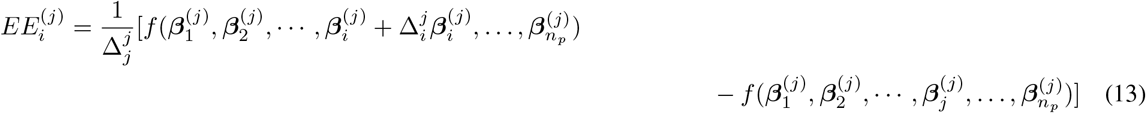

where 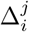 is a fixed size of variation of the input parameters and is in the range of [0, 1]. The sampling strategy involves designing *n* trajectories (also known as number of EEs) and each trajectory is composed of *n_p_* + 1 points, where the additional point refers to the perturbation of one specific parameter.

Two statistical indices based on the EEs are derived in order to evaluate the sensitivity of each parameter on the model output, namely the mean 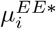 and the standard deviation 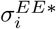, which are given as

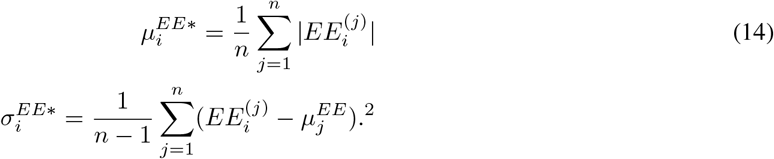

where *n* is the total number of trajectories and *i* is the *i*–th parameter analyzed. The indices 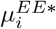 and 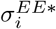 represent the influence parameter *i* has on the model out and the extent to which it nonlinearly interacts with other parameters, respectively.

The sensitivity indices are normalized as follows,

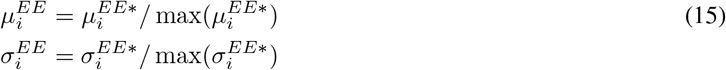

### 3.2 Sobol Method

The Sobol method is a variance-based analysis and in this approach the input parameters are considered as random vectors, characterized by a joint probability distribution *p**_β_***. With a slight abuse of notation, we represent the stochastic version of model as *y* = *f*(***β***), where ***β*** represents the input parameter vector of interest, and the Hoeffding-Sobol decomposition is given by,

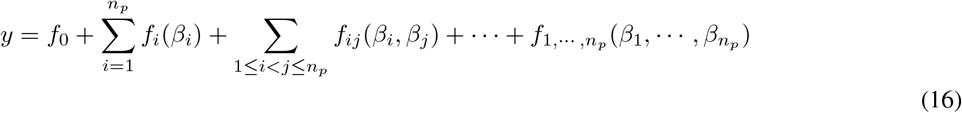

under the assumption that *p**_β_*** = *p*_*β*_1__ ⊗…⊗ *p_β_n_p___*, i.e., the input parameters are independent. Here,

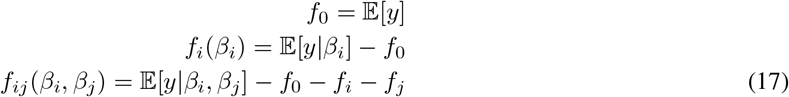

Here, the mean value is represented by *f*_0_, and the recursive definition of the conditional expectations in increasing order characterize an unique orthogonal decomposition of the model’s output [36].

The total variance of the model output can now be represented as,

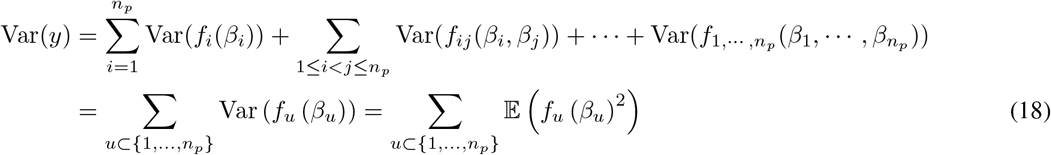

The individual impact of *β_i_* on *y* = *f*(***β***) can be measured by the first-order Sobol indices *S_i_*, defined as,

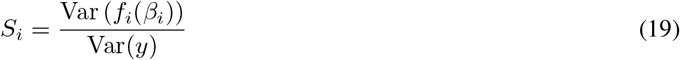

for *i* ∈ {1,… *n_p_*}.

The interaction between two input parameters is given by the second-order Sobol indices, *S_i,k_*, defined as

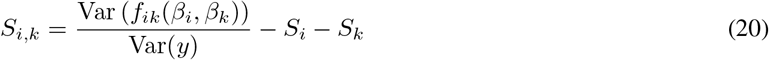

for (*i* ∈ {1,…, *n_p_*}, *k* ∈ {1,…, *n_p_*}).

Furthermore, the total-order Sobol indices that take into account the impact of the nonlinear interactions between the parameters *β*_1_,…, *β_n_p__*, which are are encoded into the functions *f_u_,* with |*u*| ≥ 2 can defined as,

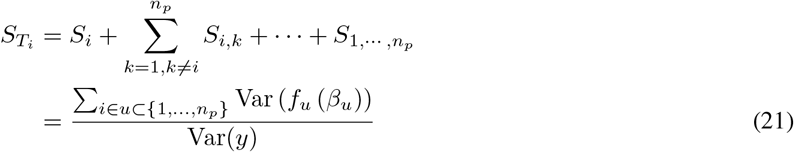

for *i* ∈ {1,… *n_p_*}.

### 3.3 Results of SA

Table 2 shows the lower and upper bound of the parameters for the Jansen-Rit NMM that was used for the Morris and Sobol method. The lower and upper bound values were based on the values reported in [37] and the references within. Since we set *A* = 6 mV and *B* = 70 mV to generate normal EEG instead of the usual *A =* 3.25 mV and *B* = 22 mV, the lower and upper bound for *A* and *B* were modified accordingly. For the Morris method, we set the the number of trajectories *n* = 1000 and 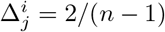. Choosing a higher number of trajectories did not qualitatively change the result. For the Sobol’s method, algorithm proposed in [20] was used and base sample size of 5000, 10000 and 15000 samples was tried. Using base sample size less than 15000 resulted in negative sobol indices indicating non-convergence and large approximation error [24]. Thus base sample size was set to 15000 samples for Sobol’s analysis.

**Table 2:** Lower and upper bound of the Jansen-Rit NMM parameters for the Morris and Sobol method

| Parameter | Lower bound | Upper bound |
| --- | --- | --- |
| $A$ | 6 | 10 |
| $B$ | 30 | 70 |
| $a$ | 10 | 150 |
| $b$ | 10 | 150 |
| $C_1, C_2$ | 50 | 600 |
| $C_3, C_4$ | 30 | 600 |
| $v_0$ | 5 | 7 |
| $e_0$ | 2 | 3 |
| $r$ | 0.5 | 0.6 |
| $\mu$ | 0 | 200 |
| $\sigma$ | 1 | 1000 |

For the Morris method, we recall that a high value for the normalized mean 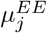 indicates that a parameter has great influence on the model output and a high value for the normalized standard deviation 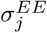 indicates that the parameter has considerable nonlinear interactions with other parameters. In the case of Sobol indices, higher values for the first-order index *S_j_* indicate that the parameter has a great influence on the model output and higher values for the total-order index *S_T_j__* indicates nonlinear interactions with other parameters. Consequently, the quantity *S_T_j__* – *S_j_* indicates how much the *j*–the parameter is involved in interaction with other parameters.

Since the NMM output is a function of time, all the sensitivity indices were computed for each time-point and then averaged over all the time points (*K* = 100000 samples). The NMM equations were solved using Runge-Kutta method with a step-size of *dt* = 0.001.

Figure 1 shows the sensitivity indices 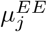 and 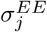 obtained with Morris method for the NMM parameters. It can be seen that parameters *C*_4_ has the highest values for both 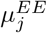 and 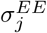 followed by *a.* Parameters *A* and *B* show similar values for 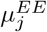 but parameter *B* has much lower value for 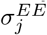 compared to *A*. Parameter *b* also shows high value for 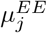 with low value for 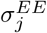. All other parameters show significantly lower values for 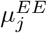, which relate to the direct impact the parameters have on model output, compared to the 5 parameters – *C*_4_, *a, b, A* and *B*. Also, among these 5 parameters, parameter *b* and *B* show the lowest values for 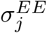 (*B* followed by *b*), which relates to the nonlinear interaction the parameter has with other parameters.

**Figure 1:**
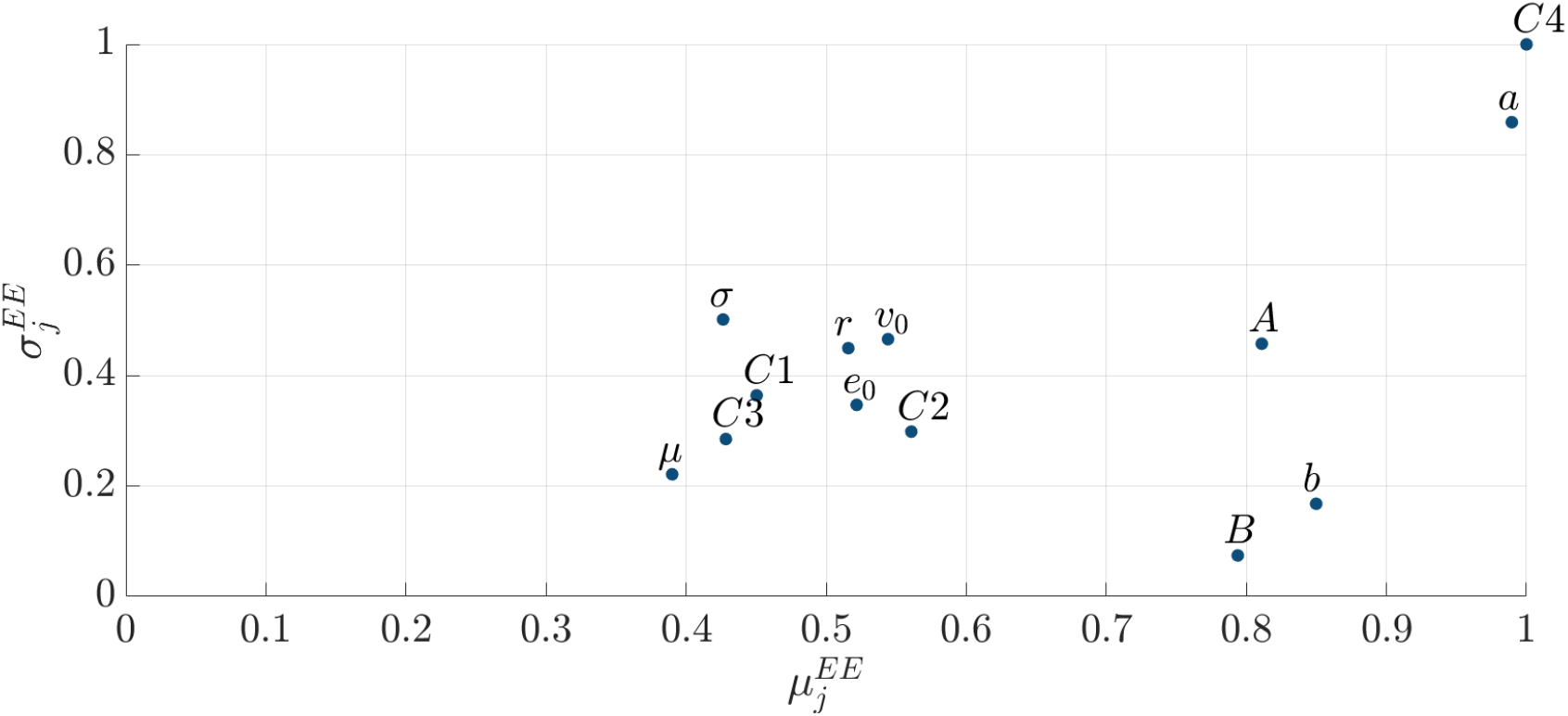
Normalized SA results for the 13 parameters of JR-NMM using the Morris method showing sensitivity indices mean 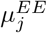 on the x-axis and and standard deviation 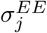 on the y-axis.

Figure 2 (a) shows the first-order index *S_j_* and total-order index *S_T_j__* using Sobol method for the NMM parameters. Parameter *C*_4_ has the highest *S_T_j__* followed by *a*. Parameter *C*_4_ also has the highest *S_j_*, followed by parameter *B*. Apart from parameters *B, a, b* and *C*_4_ other NMM parameters show negligible values for the first-order index *S_j_*, which relates to the direct influence of the parameter on model output. Among these parameters – *B, a, b* and *C*_4_, parameters *B* and *b* have the lowest values for *S_T_j__* (*B* followed by *b*). Figure 2 (b) shows the quantity *S_T_j__* – *S_j_*, which can be considered as a measure of a parameter’s contribution to all the interactions with other parameters. Parameters *B* and *b* again show low values for *S_T_j__* – *S_j_* compared to *C*_4_ and *a*. Note that we computed the quantity *S_T_j__* – *S_j_* for *C*_4_, *a, b* and *B* as only these parameters have considerable impact (based on *Sj*) on the model output.

**Figure 2:**
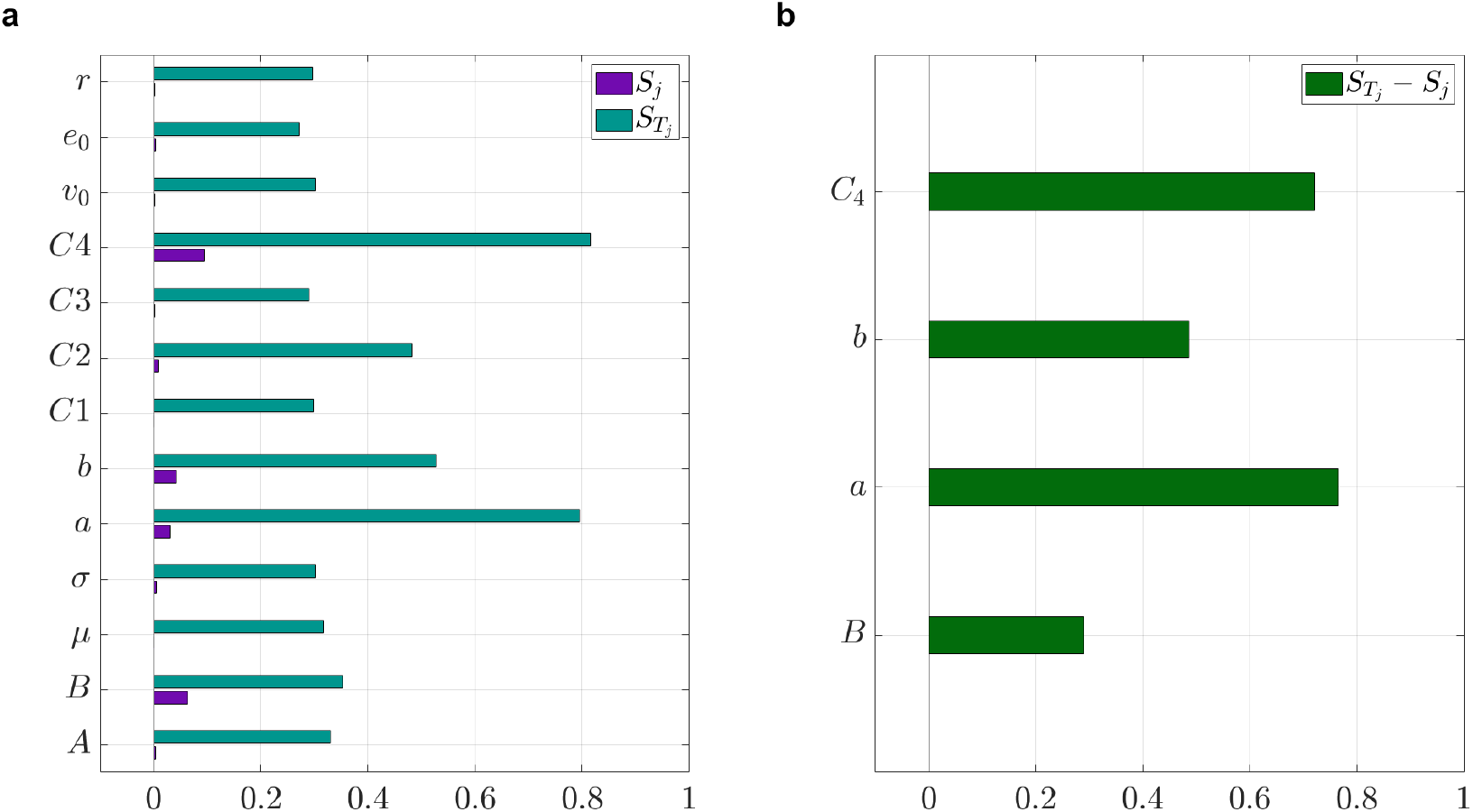
SA results for the 13 parameters of JR-NMM using the Sobol method showing (a) First-order index *S_j_* and total-order index *S_T_* and (b) The difference *S_T_* – *S_j_*

Overall, results from both Morris method and Sobol indices show that parameters *B* and *b* have considerable impact on the NMM model output and also show little interaction with other parameters. In contrast, although parameters *a* and *C*_4_ (based on Sobol and Morris method), and parameter *A* (based on Morris method) also have considerable impact on model output, they interact nonlinearly with other parameters to a great extent. Other parameters have negligible influence on the NMM output.

## 4 Joint estimation of states and parameters of NMM with UKS-EM

We propose an EM algorithm combined with UKS, hereafter known as the UKS-EM algorithm to jointly estimate the state and parameters of the Jansen-Rit NMM. Based on the results from SA shown in Section 3.3, the parameters *b* and *B* have the most influence on model output with considerably little interaction with other parameters and will be estimated with the UKS-EM algorithm, with other NMM parameters set to standard values as shown in Table 1. In addition the parameters, the UKS-EM algorithm will also estimate the states **z***_k_* (see equation 11).

The EM algorithm is a popular approach to estimate the unknown state and parameters, given the observation data. By combing the unobserved state and parameter as 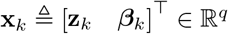 where 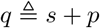, the state-space model in in Equation 10 can be re-written as,

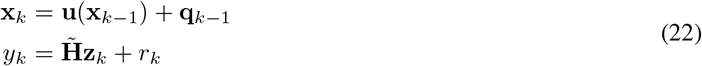

where 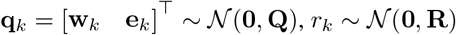 and,

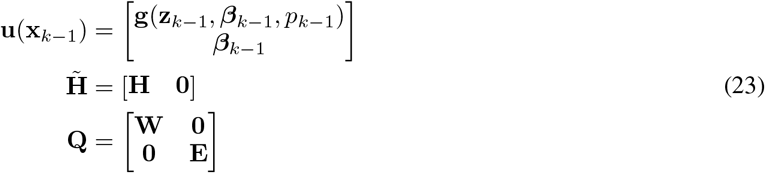

The task is to now infer the unknown state **x***_k_* and parameter ***θ*** = [**Q**, **R**], given the observation **y***_k_*. Let **X** = **x**_1:*N*_ = {**x**_1_,…, **X***_N_*} and **Y** = **y**_1:*N*_ = {**y**_1_,..**y**_*N*_}. From Bayes’ rule ^1^,

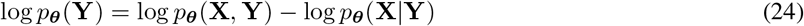

The expectation maximization (EM) algorithm aims to maximize the lower bound of the marginal log-likelihood, log *p_**θ**_*(**Y**), by maximizing the expected value of the complete data log-likelihood,

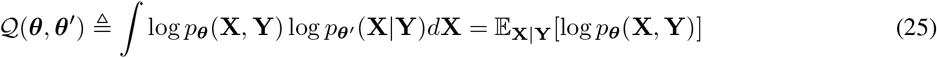

in an iterative manner, where ***θ**′* is the current estimate of ***θ***. Starting with an initial value ***θ***_0_, the EM algorithm iterates between the two steps until convergence or the maximum number of iterations is reached:

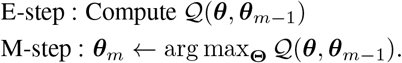

In addition to the estimate of the parameter ***θ**_m_,* the full posterior distribution *p_θ_m__*(**X**|**Y**) is also obtained at every iteration *m* of the EM algorithm, from which the unobserved states **X** can be inferred.

Due to the Markovian structure, we can write the complete data log-likelihood as follows,

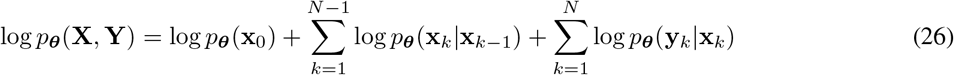

Taking the expectation of the above quantities, we can write

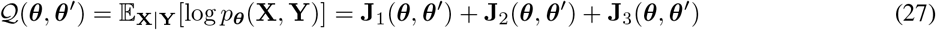

where,

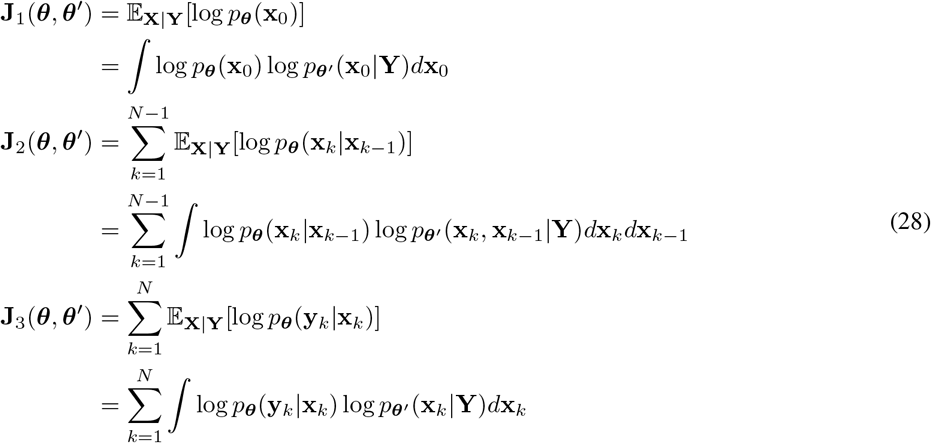

To evaluate these expectations, we need the smoothing distributions log*p_**θ**_* (**x***_k_*|**Y**) and log*p_**θ**_* (**x**_*k*+1_, **x***_k_*|**Y**). In this work we use the UKF and UKS to approximate these expectations.

### 4.1 Expectation step

#### 4.1.1 The unscented Kalman filter (UKF)

Let 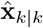 and **P***_k|k_* represent the mean and covariance of the augmented state 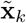. We now define the following sigma points,

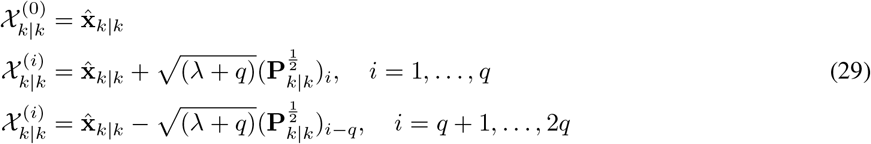

where *λ* = *α*^2^(*q + c*) – *q* is a scaling parameter and the choice of *α* influences the spread of the sigma points around the mean 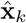. The subscript *i* in 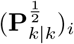 represents the *i*-th column of the matrix. The matrix square root 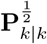 can be obtained for example using Cholesky decomposition. The weights of the sigma points for the mean terms are given as,

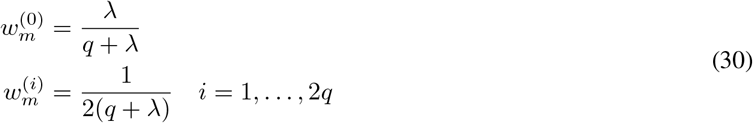

and the weights for the covariance terms are given as,

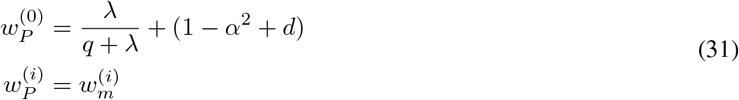

After propagating the sigma points through the nonlinear function **f**, we can compute the mean 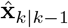 and covariance **P**_*k*|*k*-1_ of the predictive distribution *p*(**X***_k_*|**y**_1:*k*-1_) as,

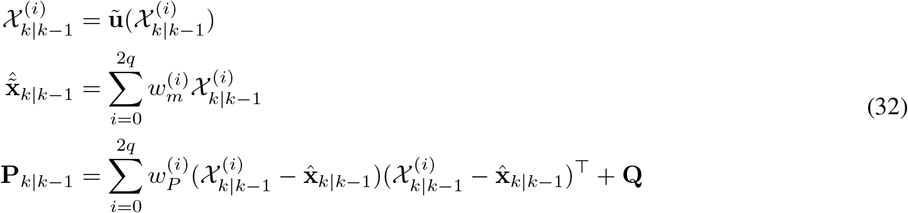

In the measurement update step, a new set of sigma points are formed using 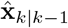 and **P**_*k*|*k*-1_,

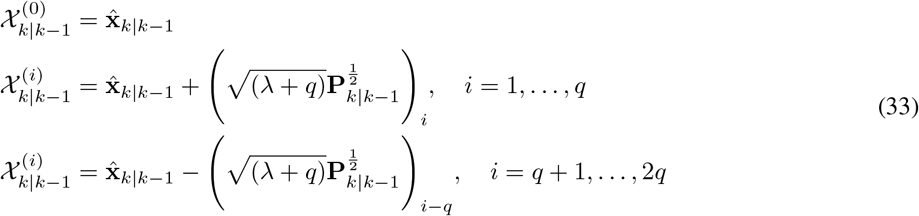

which is transformed to the measurement space through the linear transformation in case of the model we are using (see equation XX),

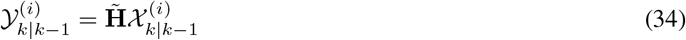

to compute the mean 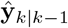 and covariance **M**_*k|k*-1_ of the measurement predictive distribution *p*(**y***_k_*|**y**_1:*k*-1_) and the cross covariance **Z**_*k|k*-1_ of the state and measurement predictive distributions,

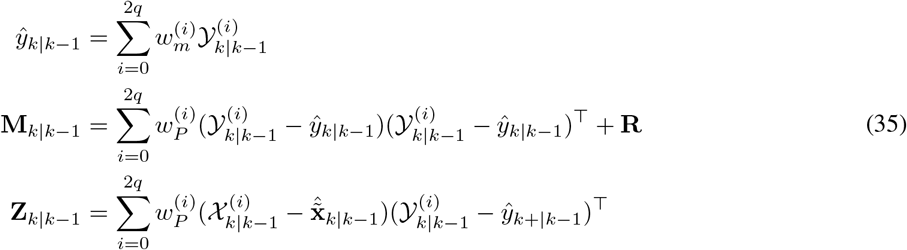

The filtered mean and covariance of the augmented state is then given by the following equations,

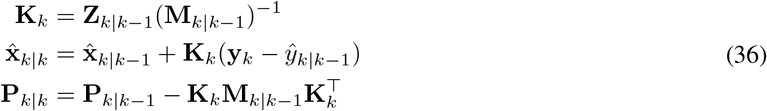

where **K***_k_* is the Kalman gain.

#### 4.1.2 The unscented Kalman smoother (UKS)

The unscented Rauch–Tung–Striebel smoother (also known as UKS) [34] is used to compute the smoother mean 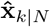 and covariance **P***_k|N_* as,

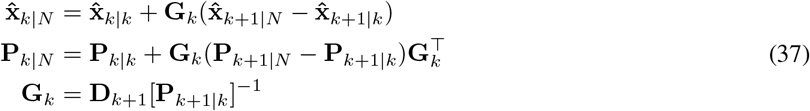

where 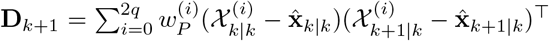. Note that in order to compute **J**_2_(***θ, θ’***), we need the pairwise smoothing distribution *p_**θ**_* (**x***_k_*, **x**_*k*-1_|**Y**), which is given as

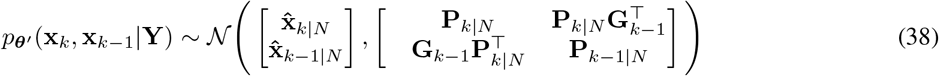

Expanding the quantities within the expectation operator 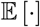 in Equations 27 and 28, we have

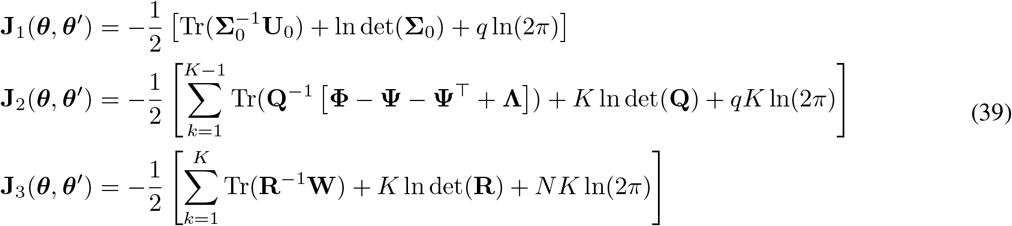

where,

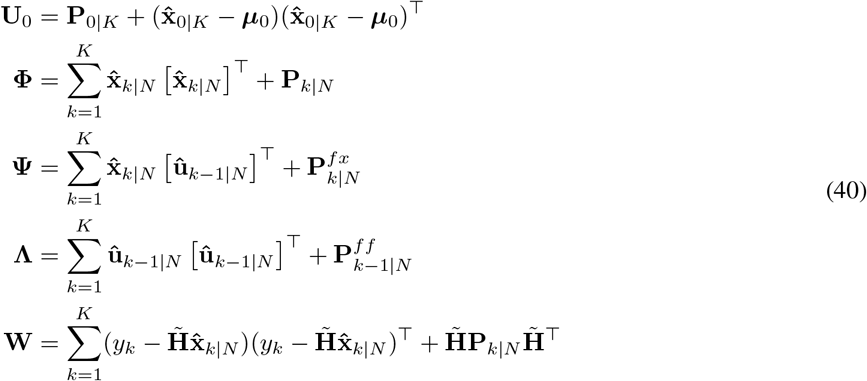

The covariance **P***_k|N_* is given by the Kalman smoother (see Equation 37). We define the following sigma-points,

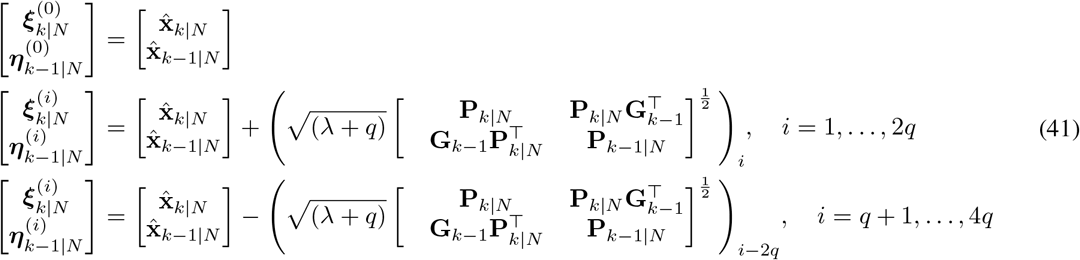

in order to compute the covariances 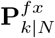 and 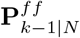 and the mean 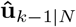,

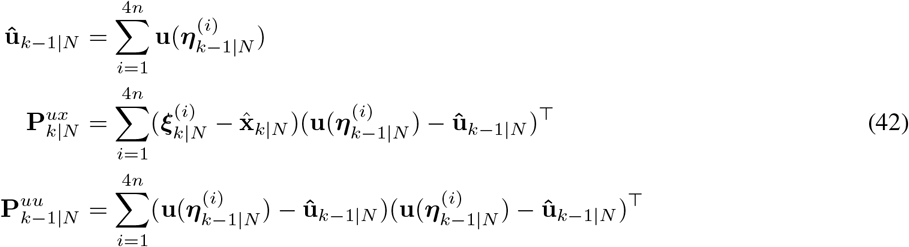

### 4.2 Maximization step

In the maximization step, the covariance matrices **Q** and **R** are estimated by maximizing the terms **J**_2_(***θ, θ’***) and **J**_3_(***θ, θ’***) w.r.t **Q** and **R** respectively.

#### 4.2.1 Update of Q

At iteration *m*, to maximize **J**_2_(***θ, θ’***) w.r.t **Q**, we set 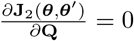 we get [38],

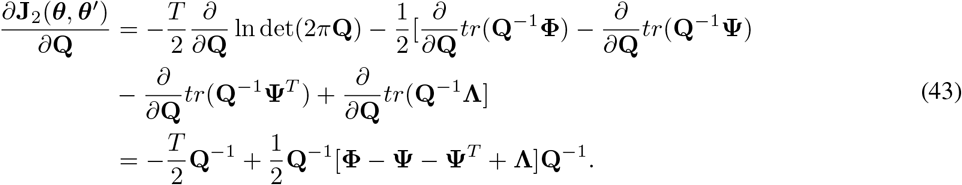

Setting the above derivative to zero, we obtain 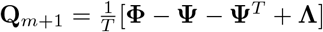.

#### 4.2.2 Update of R

At iteration *m*, setting 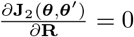 we get [38]

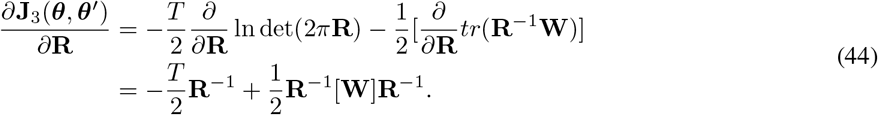

Setting the above derivative to zero, we obtain **R**_*m*+1_ = **W***/T*.

## 5 Simulations

Based on the results from SA (See Section 3.3 and Figures 1 and 2), we simulated EEG signals using JR NMM by varying the parameters *B* and *b* and keeping all other parameters fixed to the standard values shown in Table 1 [5].

Figure 3 shows the simulated EEG signal of duration *T* = 100 seconds, obtained using Jansen-Rit NMM output when parameters *B* and *b* are varied to produce normal EEG followed by rhythmic discharge of spikes. With *B* = 70 and *b* = 50, the NMM produced normal EEG as its output. But as *B* is reduced to 50 at *T* = 20 seconds rhythmic discharge of spikes appear, which increases in frequency as *b* is increased to 100 at *T* = 60 seconds.

**Figure 3:**
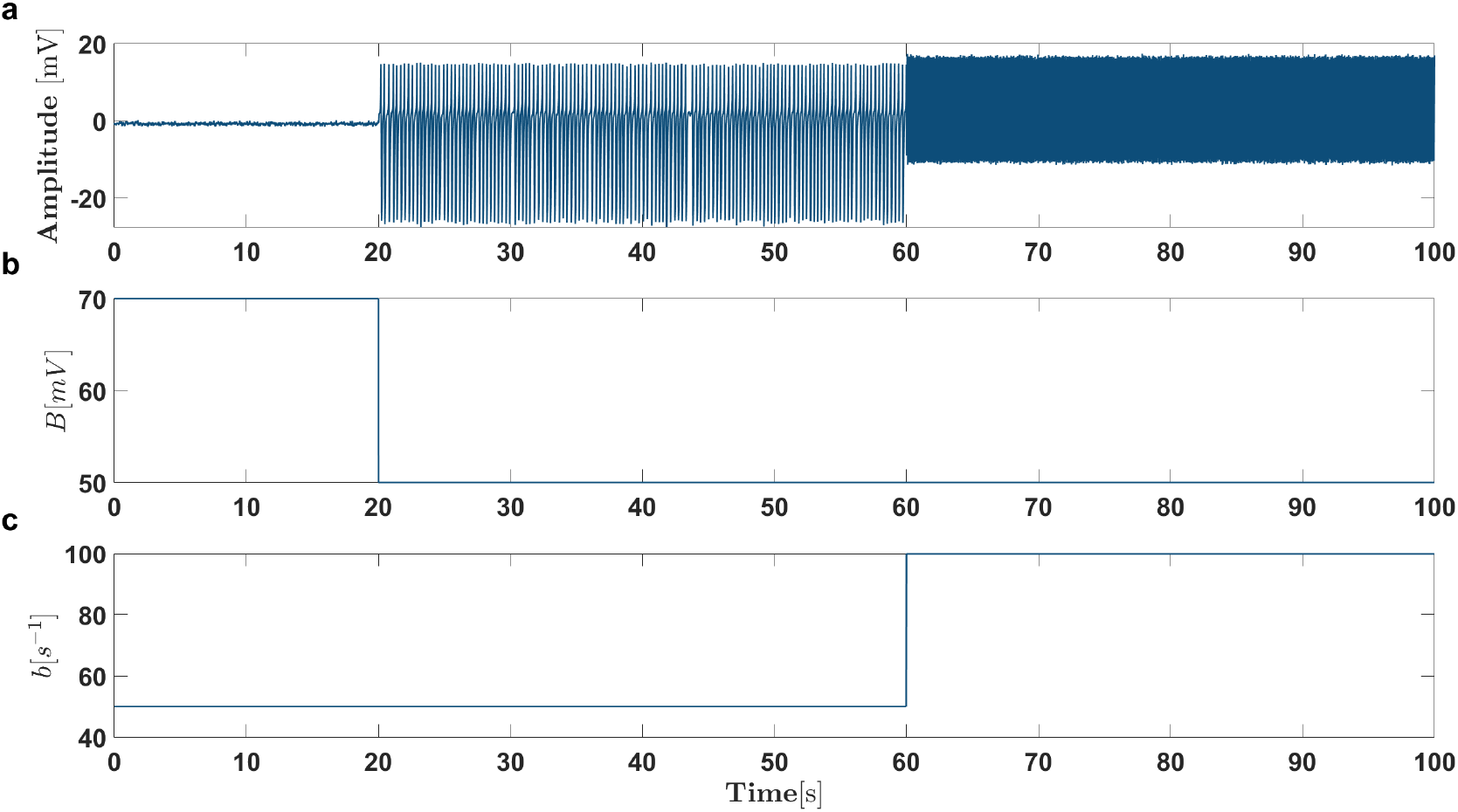
Simulation of EEG using JR NMM. Based on the results of SA, parameters *B* and *b* are identified as the most sensitive parameters. Starting with the nominal value of *B* = 70 mV and *b* = 100 s^-^1 which produces normal EEG, reducing *B* to 50 mV produces rhythmic activity [5]. Increasing *b* to 100 s^-^1 increases the frequency of the rhythmic activity. All other JR NMM parameters are fixed to the nominal values shown in Table 1.

In order to asses the performance of parameter estimation with UKS-EM algorithm under the influence of noise, varying levels of measurement noise was added (at *SNR* 1, 3, 5 and 10) to the simulated EEG shown in Figure 3 according to,

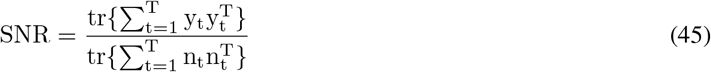

where 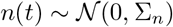 with 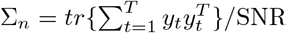, is the simulated noise signal at the specified SNR and *y*(*t*) is the noise-free output of the NMM. In Figure 4 simulated EEG at various SNRs is shown.

**Figure 4:**
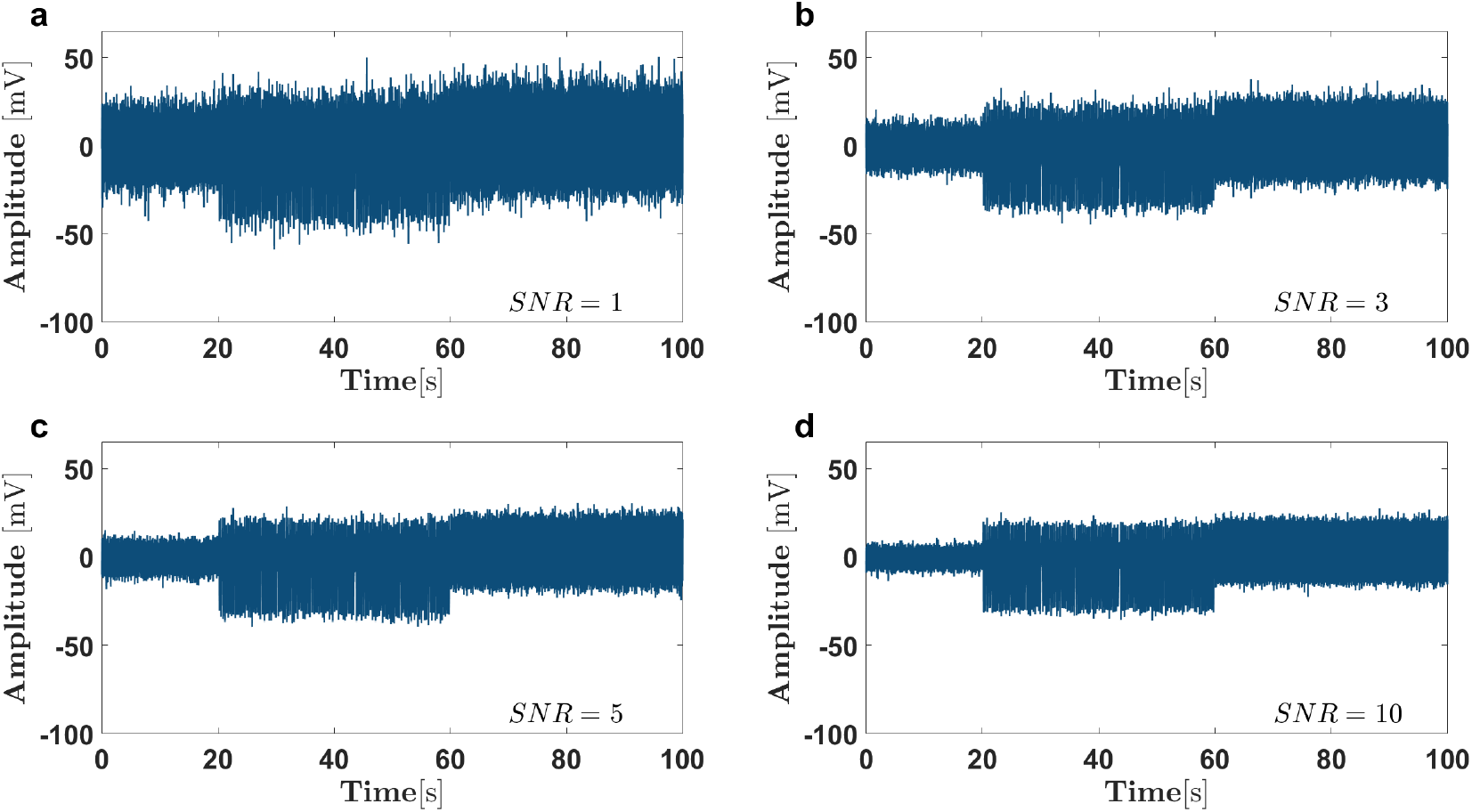
Output of the JR NMM for varying levels of noise with *SNR* ranging from 1 to 10.

### 5.1 Results

Figure 5 shows the parameter estimation results at various SNRs for parameters *B* and *b,* using the UKS-EM algorithm for the simulated EEG signals, with the shaded region representing confidence intervals (CI) corresponding to two standard deviations (2*σ*), which is obtained from the Kalman smoother covariance. The UKS-EM algorithm provided more accurate estimate of the parameter *b* with smaller CI across the range of SNRs compared to parameter *B*. The UKS-EM estimates of parameter *B* improved with CI becoming smaller as SNR increased. Table 3 shows the root mean squared error (RMSE) for parameters *B* and *b* at various SNRs. The RMSE for parameter *b* remains low even at high SNRs, while RMSE of parameter *B* improves with increasing *SNR.*

**Figure 5:**
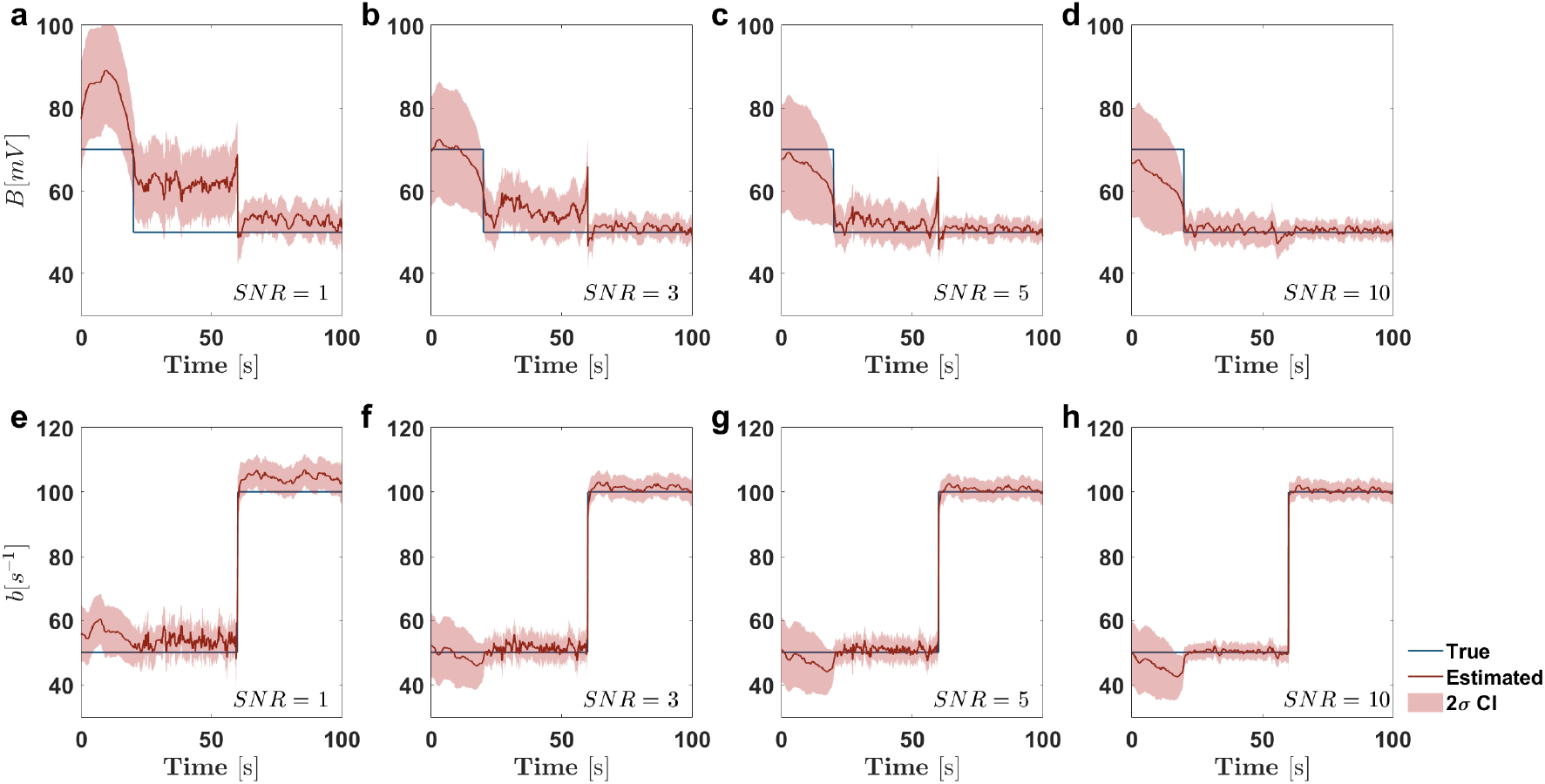
Estimation of the NMM parameter *B* and *b* by UKS-EM algorithm for a range of SNRs. The shaded region represents the 2*σ* CI.

**Table 3:** RMSE at various SNRs for NMM parameters *B* and *b*.

| NMM Parameter | RMSE |  |  |  |
| --- | --- | --- | --- | --- |
|  | SNR = 1 | SNR = 3 | SNR = 5 | SNR = 10 |
| $B$ | 10.2 | 3.96 | 3.11 | 3.54 |
| $b$ | 4.2 | 1.95 | 1.91 | 1.89 |

## 6 Real data

To validate our methods on real data, we used intracranial EEG (iEEG) data of epileptic patients recorded and analyzed at the Sleep-Wake-Epilepsy-Center (SWEC) of the University Department of Neurology at the Inselspital Bern and the Integrated Systems Laboratory at ETH Zurich. The data is freely available at http://ieeg-swez.ethz.ch/. Briefly, the database consisted of a total of 100 iEEG datasets recorded from 16 patients using grid, strip and depth electrodes. The iEEG signals were acquired at a sampling frequency of 512 Hz. A fourth-order Butterworth filter between 0.5 and 150 Hz was used to bandpass filter the data. In each of the iEEG recordings, the pre-seizure and post-seizure duration were 3 minutes and the seizure duration itself ranged from 10 to 1000 seconds. All the iEEG recordings were visually inspected by an EEG board-certified experienced epileptologist for identification of seizure onsets and endings and exclusion of channels continuously corrupted by artifacts.

In order to reduce the computational cost, for the purpose of our analysis, we further segmented the iEEG recordings to include just 1 minute of data before and after the seizure and selected one iEEG recording per patient. Figure 6 shows the iEEG recording from an exemplary patient.

**Figure 6:**
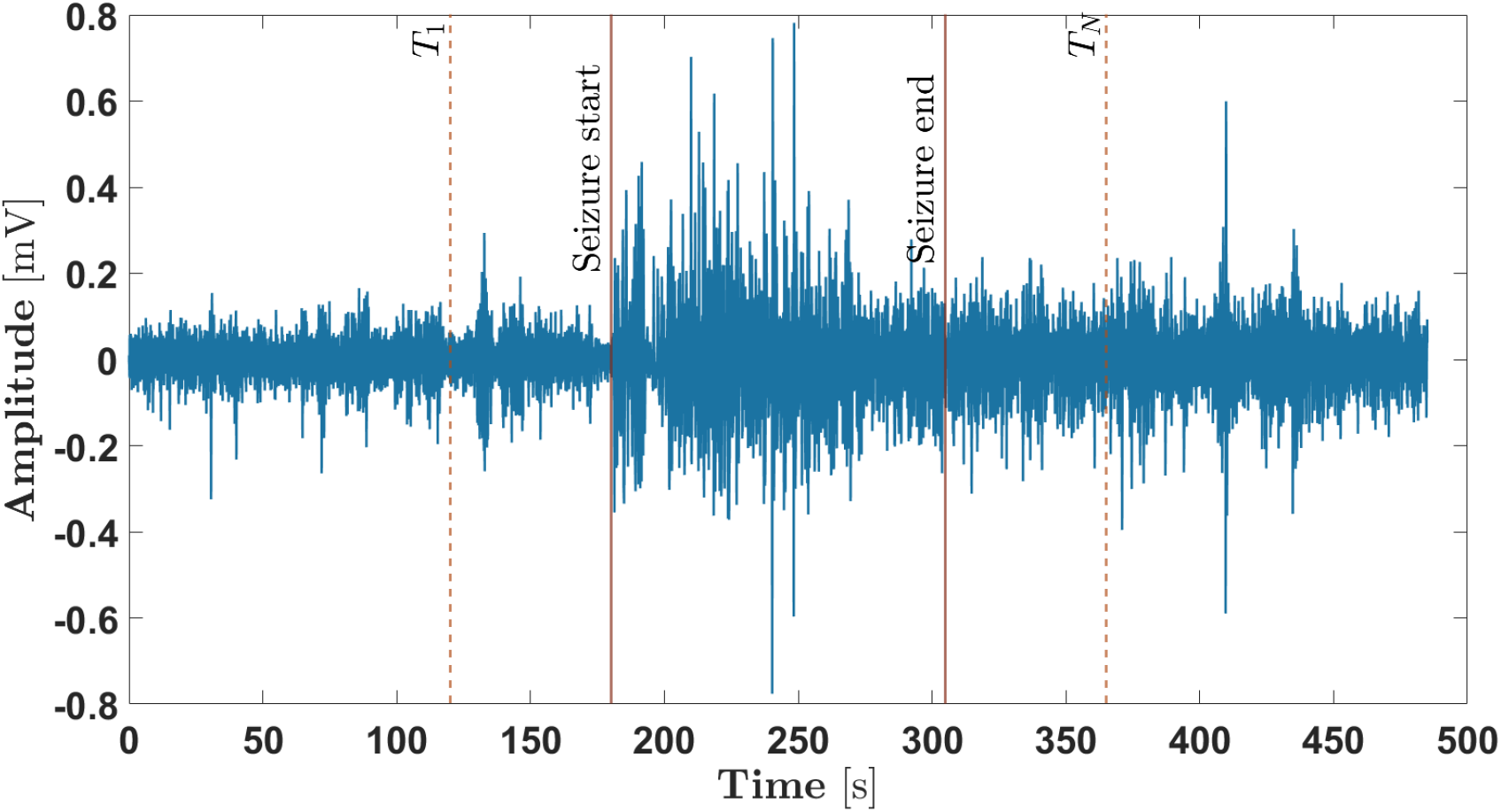
An example of one-channel iEEG recording from an epileptic patient obtained from the open database (ref). The start and end of the seizure is shown with solid red vertical lines. For the purpose of this work, iEEG data one-minute before and one-minute after the seizure, shown with dotted red lines (*T*_1_ to *T_N_*) was considered.

### 6.1 Results

In Figure 7, the results of tracking the NMM parameters *B* and *b* by the UKS-EM algorithm is shown for an exemplary subject. We can see from Figure 7 that both the parameters *B* and *b* remain near their nominal values of 70mV and 50s^-1^ during the pre-seizure period. At around 60 seconds, which marks the start of the seizure, parameter *b* increases followed by a drop in parameter *B.* During the seizure period, the value of parameter *B* remains under 70mV. In contrast parameter *b* mostly remains near its nominal value during the seizure period but is characterized by periods of transient increase, which corresponds with increase in the frequency of ictal activity during seizure period. During these brief periods, parameter *B* is characterized by further drop in its value. After the seizure, both the parameters *B* and *b* return near their nominal values.

**Figure 7:**
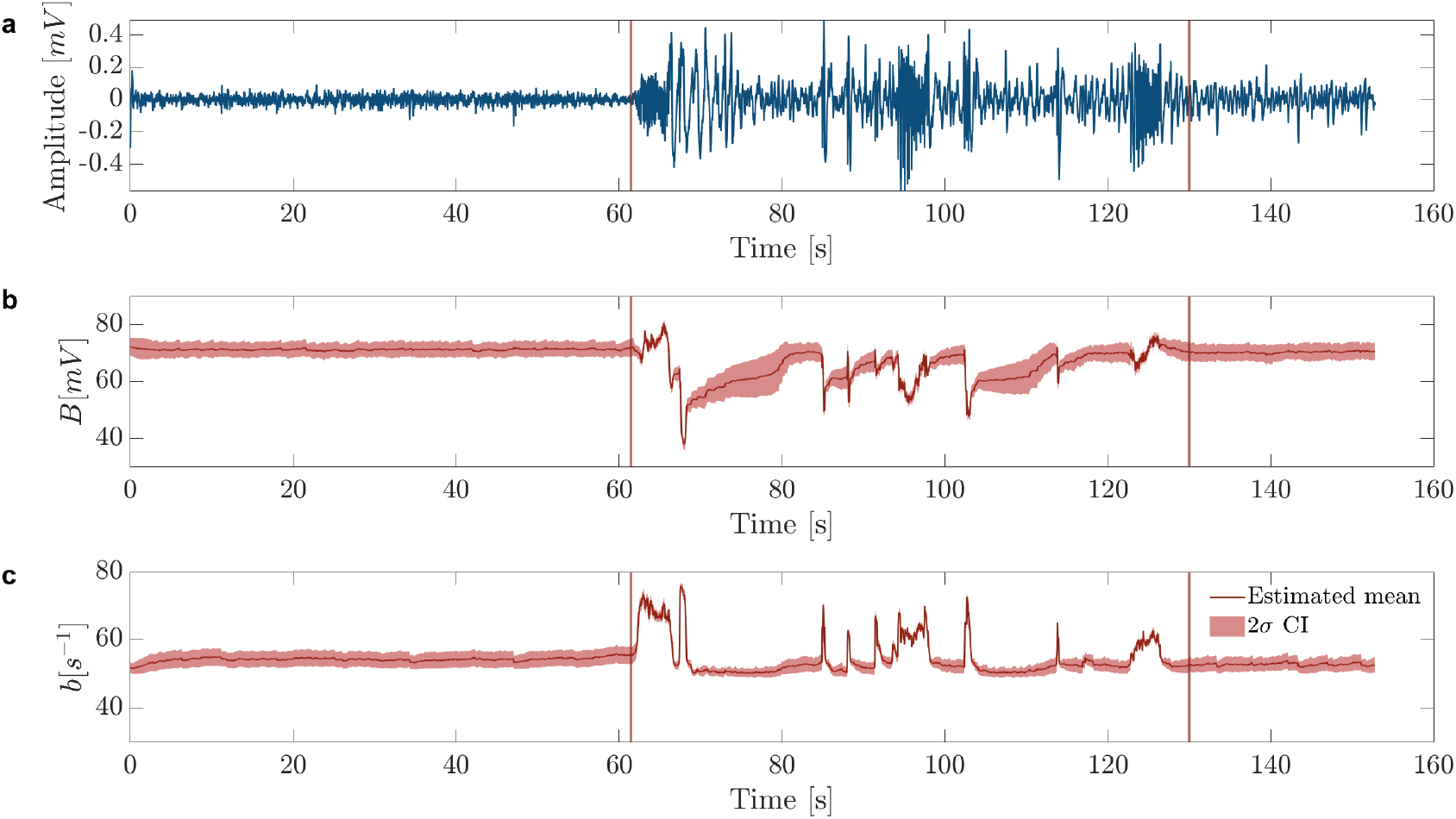
Parameter estimation results from UKS-EM algorithm. (a) Exemplary iEEG data from one epileptic patient. Estimated parameter mean from UKS-EM for (b) Parameter *B* and (c) Parameter *b*. The shaded region in (b) and (c) represents CI corresponding to two standard deviations.

Figure 8 shows the mean ± standard deviation of parameters *B* and *b* for the pre-seizure, seizure and post-seizure period for all the sixteen subjects. We can see that for all the subjects parameter *B* tends to have lower values during the seizure period compared to pre-seizure and post-seizure period. In contrast, parameter *b* has higher value during the seizure period compared to pre-seizure and post-seizure period for all the subjects. For all the subjects we found statistically significant (*p* < 0.01) differences for *B* and *b* between the pre-seizure and seizure, seizure and post-seizure period.

**Figure 8:**
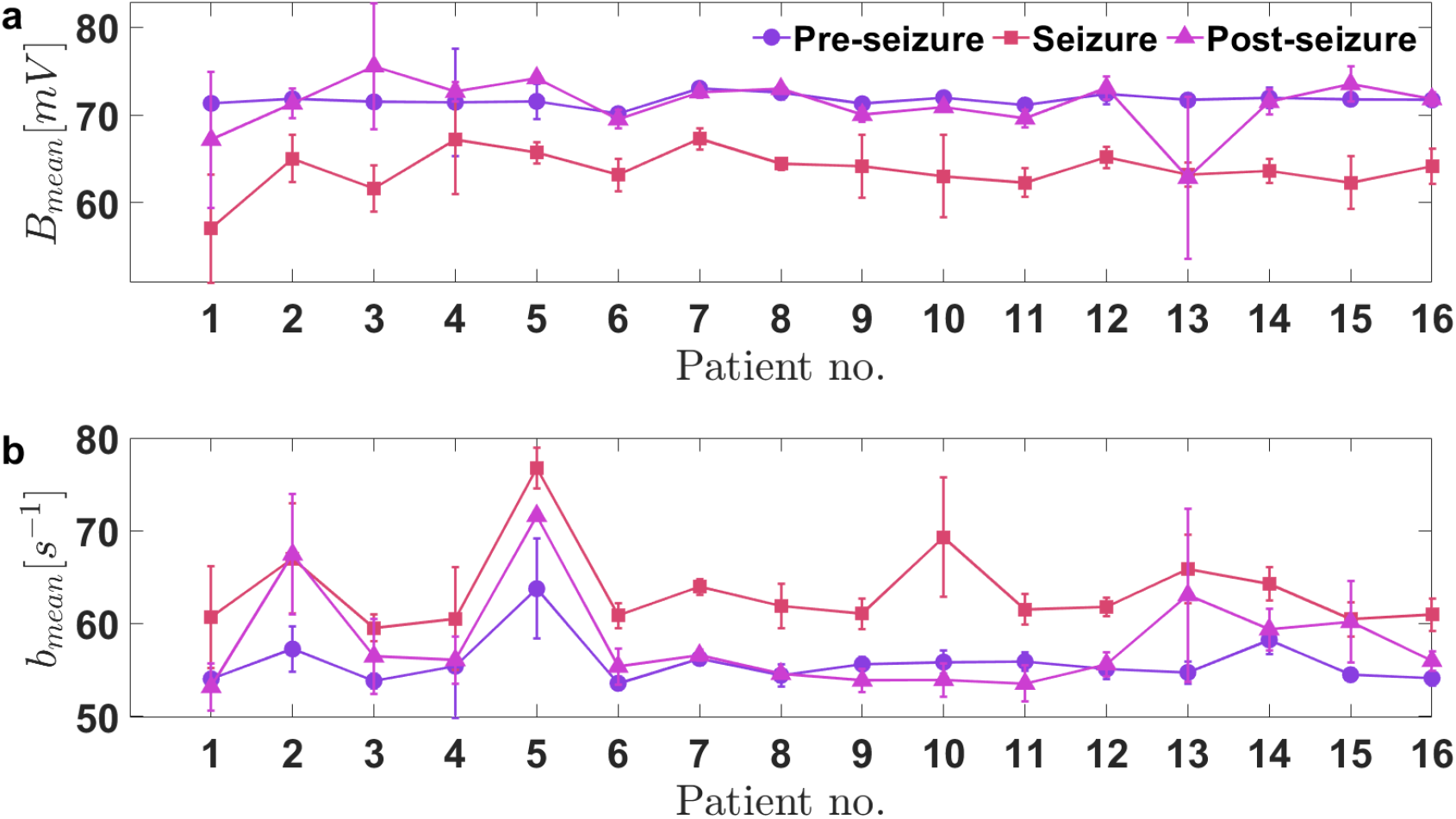
Mean shown with solid lines and standard deviation shown with error bars for parameters *B* and *b* during pre-seizure, seizure and post-seizure for 16 epileptic patients.

In Figure 9 we show the mean values of parameters *B* and *b* during the pre-seizure, seizure and post-seizure period, averaged over 16 subjects, with error bars representing the standard deviation. We used a two-tailed t-test to asses the significance in paramter changes between the three different periods – pre-seizure, seizure and post-seizure. The changes in parameters *B* and *b* were statistically significant at the group level between pre-seizure and seizure period (*p* < 0.001) and also between seizure and post-seizure period (*p* < 0.001 for *B* and *p* < 0.01 for b), while the changes in parameters were not found to be significant between pre-seizure and post-seizure periods.

**Figure 9:**
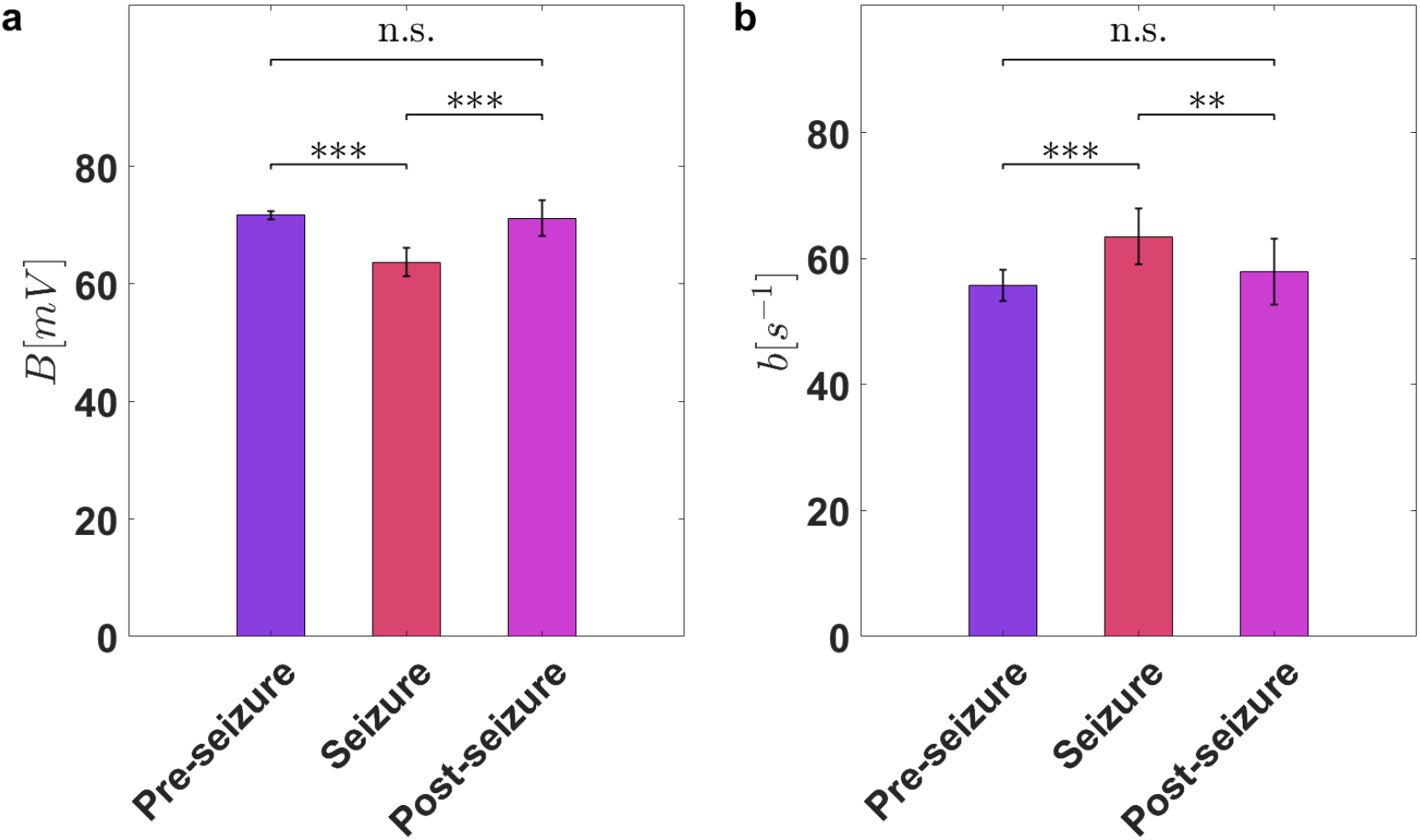
Comparison of UKS-EM estimation results at the group level (*N* = 16 patients) between pre-seizure, seizure and post-seizure periods for (a) Parameter *B* and (b) Parameter *b*. The mean values of parameters *B* and *b* during the pre-seizure, seizure and post-seizure period, averaged over 16 patients is shown with the error bars representing the standard deviation. \**p* < 0.05, \*\**p* < 0.01, \*\*\**p* < 0.001 and n.s is not significant,.

## 7 Discussion

In this study, we introduced global SA techniques namely the Morris method and Sobol method to investigate the influence of NMM parameters on the model output and the extent of interaction between the NMM parameters. We used the JR model as an example of NMM. The goal of such an investigation was to identify a set of NMM parameters that can be reliably estimated from EEG data. Furthermore, for parameter estimation from EEG data, we implemented the UKS-EM approach that combines the UKS with the EM algorithm.

Our results based on both the global SA techniques showed that the dendritic time constant of the average inhibitory PSPs, 1/*b* and the average inhibitory synaptic gain parameter *B* has considerable influence on model output along with having relatively low nonlinear interaction with other parameters, implying their reliability for parameter estimation from the EEG data. The sensitivity indices based on the mean and standard deviation from Morris method and first-order index and total-order index based on Sobol method, both indicated that other NMM parameters such as average time constant of the excitatory PSPs 1/*a*, the average number of synaptic contacts in the inhibitory loop *C*_4_, has greater influence on the model output, but also show greater extent of nonlinear interaction with other NMM parameters when compared to 1/*b* and *B*, thus questioning their reliable estimation from data. Global SA results also indicated that the excitatory gain parameter *A* has similar levels of influence on model output as the inhibitory gain parameter *B*, but parameter *A* showed higher degree of nonlinear interaction with other NMM parameters, implying that *B* can be estimated more reliably than *A*. All other NMM parameters showed lower degree of influence on the model output compared to 1/*b* and *B* and thus suggesting that these cannot be reliably estimated from EEG data.

Based on our SA results, the information on the parameter sensitivities of the JR NMM provided by the Morris method was similar to the Sobol method, but at a reduced computational cost, since the Morris method requires a total of *n*(*n_p_* + 1) evaluations, where *n_p_* is the number of parameters and *n* is the number of trajectories and the Sobol method requires n(n_p_ + 2) model evaluations [20]. Thus, the Morris method can be considered as a practical choice of SA method to analyze NMMs with large number of parameters.

Based on the global SA results, we estimated the parameters *B* and *b* using an UKS-EM algorithm. We showed using simulations that the parameter *b* can still be reliably estimated from EEG data even at high levels of measurement noise (SNR = 1) and parameter *B* can be reliably estimated under moderate levels of noise (SNR > 1). The EEG was simulated by varying both *B* and *b* to produce rhythmic discharges that gradually increases in frequency as *b* is increased. The results from UKS-EM estimation aligns well with the global SA analysis which suggested that the parameter *b* has a greater influence on output compared to parameter *B*. Specifically based on the results from Sobol indices, we showed that both *B* and *b* have similar level of interaction with other parameters, but *b* has a greater influence on model output compared to *B*. The results from the simulations using UKS-EM algorithm is in line with this observation.

Using real data in the form of iEEG data from 16 epileptic patients, we further validated our approach. Our results show statistically significant differences in the changes in both the parameters *B* and *b* between the pre-seizure and seizure and the seizure and post-seizure, both at the individual level as well as group level. Previous studies have attempted at estimating mainly the excitatory and inhibitory gain parameters, i.e., *A* and *B* respectively using epileptic EEG data. Specifically we observed that during the seizure period the value of parameter *B* drops below the pre-seizure level and the value of parameter *b* increases above the pre-seizure levels. During the post-seizure period, the values of these parameters returns back to the pre-seizrue period. This observation was consistent across all the 16 subjects. The observation on the changes in parameter *B* during the seizure period is consistent with the previous studies. Wendling and colleagues [9, 5] showed that reduction in the inhibition gain *B* leads to the transition to seizure like activity. In their model, further reduction in *B* also lead to high frequency discharge due to the presence of a fast inhibitory population with a ten times smaller time constant compared to the dendritic time constant *b*. Since we used the original JR model that has just one inhibitory population as opposed to two inhibitory population - slow and fast as proposed in Wendling and colleagues [9, 5], the high frequency activity was explained by shorter dendritic time-constant which is represented by an increase in the value of parameter *b*, which corresponded with high frequency-like activity during the seizure period as seen in Figure 7. This was also demonstrated in our simulations where the frequency of the rhythmic discharges increased as value of *b* was increased from 50 to 100 (or conversely the dendritic time-constant 1/*b* was reduced). These results provide a new intuition on how JR model can also explain high frequency-like activity without the need of an additional fast inhibitory population. The reduction in the dendritic time-constant 1/*b* produces faster IPSPs, which explains the high frequency activity and the chirping-like effect during the seizures. We believe that our results from SA which revealed that among all the JR NMM parameters, *B* and *b* can be reliably tracked, have significant implications in terms of model simplification. By also allowing the parameter *b* to vary which is normally assumed to be constant and by estimating both *B* and *b* we found that we can explain the high frequency activity in epileptic EEGs without the need for additional inhibitory populations which requires a NMM with 10 nonlinear ODEs as proposed by Wendling and colleagues [9], while JR model used here has only 6 ODEs, further reducing the computational time for Kalman filtering and smoothing for parameter estimation. Note that this is only from the point of view of model simplification for parameter estimation. There is still clearly a need to develop more physiologically plausible models such as the one proposed by Wendling and colleagues [9], Zavaglia and colleagues [39] or including astrocytic contributions as proposed in [40] for gaining deeper insights into the mechanism of epilepsy.

To the best of our knowledge, Ferrat and colleagues proposed a study to investigate the contribution of different parameters towards NMM dynamics (Wendling NMM [9]), using a random forest approach to explore the whole parameter space and determining the importance of each parameter with variable importance measure [37]. While Ferrat and colleagues focused on determining the impact of each parameter on NMM dynamics, our approach also provides the information on how much each parameter interacts with other parameters, in addition to providing the impact of each parameter on the NMM output. Furthermore, we have demonstrated the application of our analysis on real EEG data containing seizures and applied Bayesian filtering approach to estimate the sensitive parameters, while in [37] only simulations were considered.

Although, Ferrat and colleagues used the Wendling NMM, our results support their conclusion that parameter *B* and *b* have the most important role in the transition towards seizure dynamics, as seen in our results on SA and application to real iEEG data containing seizures. Interestingly, they found that the parameters of the fast inhibitory loop (the time constant g and gain G in the Wendling model), had very little impact towards transition to seizure dynamics, which relate to our previous discussion on how high-frequency activity can be generated by varying parameter *b*.

### 7.1 Limitations and future work

For the inference of the sensitive NMM parameters we used the UKS-EM approach, which requires many computations and iterations and thus is not suitable for online estimation. Adaptive Kalman filtering techniques that employ self-tuning [41] could be explored to solve the problem of unknown noise covariance matrices which we have solved through an iterative EM-like approach. Such an approach could enable online and/or faster estimation of NMM parameters.

It has been shown that for biological models the parameter ranges should be biologically feasible so that the sensitivity indices are not biased by implausible model realisations and changing the parameter range may affect the sensitivity results and thus insensitive parameters may appear sensitive, or vice versa [24]. Since NMMs are phenomenological models, the possible parameter ranges are based on previously used values that have been set arbitrarily or through fitting experimental data. In our study, we carefully set the range based on previous literature and simulations avoiding implausible model output. This is feasible with JR NMM which has only 13 parameters. For performing sensitivity analysis on NMM with larger parameters for example as the one proposed in [40] which also includes astrocytic parameters, identifying feasible parameter ranges may be more challenging. In future, more experimental studies might provide a better into the realistic range that NMM parameters could take.

Furthermore, contrary to the results reported in Ferrat and colleagues [37] where only parameters *B* and *b* had significant impact, our results showed that changes in parameters *C*_4_, *a* and *A* also had significant impact on model output. But since our results from SA also showed they had high interactions with other NMM parameters (see Figures 1 and 2) compared to *B* and *b*, we chose to estimate only *B* and *b* from the EEG data in this study. Future studies will involve also estimating second-order Sobol indices, which will provide information on which combination of parameters among *C*_4_, *a, A, B* and *b* are highly correlated.

And finally, although we have focused on epilepsy, other application areas include tracking the depth of anesthesia [18], burst suppression [42] and Parkinson’s disease [43] for which NMMs have been proposed.

## 8 Conclusion

To conclude, our work is the first attempt to perform sensitivity analysis guided Bayesian inference for the NMM parameters. Through such analysis we can set the insensitive NMM parameters to constant value and make parameter tracking more feasible by estimating the most sensitive parameters, that also interact the least with other parameters. Our results show that the average inhibitory synaptic gain *B* and the average time-constant of the inhibitory PSPs are the most important NMM parameters for reliable estimation as they have considerable impact on NMM model output and also interact less with other NMM parameters. We also proposed UKS-EM algorithm to perform parameter tracking with EEG data, demonstrated with iEEG data from 16 epileptic patients that changes in parameters *B* and *b* explain the transition towards epileptic seizures, with increase in *b* correlating with high frequency-like activity.

## 9 Acknowledgements

This project has received funding from the European Union’s Horizon 2020 research and innovation programme FETPROACT-01-2018 (RIA) awarded to the project Hybrid Enhanced Regenerative Medicine Systems (HERMES) under grant agreement No 824164.

## 10 Code and data availability

All the codes related to our work are available at https://gitlab.com/narayanps/sa-param-est.git. The sensitivity analysis was performed using the SAFE toolbox (https://safetoolbox.github.io/). The iEEG data used in this work is available at http://ieeg-swez.ethz.ch/.

## Footnotes

1 From Bayes’ rule we have 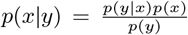. Thus, 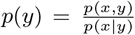. Applying log on both sides, we have log*p*(*y*) = log *p*(*x, y*) – *p*(*x|y*).

